# Intracortical dynamics underlying repetitive stimulation predicts changes in network connectivity

**DOI:** 10.1101/548180

**Authors:** Yuhao Huang, Boglárka Hajnal, László Entz, Dániel Fabó, Jose L. Herrero, Ashesh D. Mehta, Corey J. Keller

## Abstract

Targeted stimulation can be used to modulate the activity of brain networks. Previously we demonstrated that direct electrical stimulation produces predictable post-stimulation changes in brain excitability. However, understanding the neural dynamics *during* stimulation and its relationship to post-stimulation effects is limited but critical for treatment optimization. Here, we applied 10Hz direct electrical stimulation across several cortical regions in 14 patients implanted with intracranial electrodes for seizure monitoring. The stimulation train was characterized by a consistent increase in high gamma (70-170Hz) power. Immediately post-train, low-frequency (1-8Hz) power increased, resulting in an evoked response that was highly correlated with the neural response during stimulation. Using two measures of network connectivity, cortico-cortical evoked potentials (indexing effective connectivity) and theta coherence (indexing functional connectivity), we found a stronger response to stimulation in regions that were highly connected to the stimulation site. In these regions, repeated cycles of stimulation trains and rest progressively altered the stimulation response. Finally, after just 2 minutes (10%) of repetitive stimulation, we were able to predict post-stimulation connectivity changes with high discriminability. Taken together, this work reveals a relationship between stimulation dynamics and post-stimulation connectivity changes in humans. Thus, measuring neural activity during stimulation can inform future plasticity-inducing protocols.

## Introduction

Brain stimulation treatments including repetitive transcranial magnetic stimulation (rTMS) have recently emerged as clinically-effective alternatives to medications for neuropsychiatric disorders. Although rTMS is FDA-approved for certain disorders (major depression, obsessive compulsive disorder) and there are multiple ongoing clinical trials for other disorders (post-traumatic stress disorder, substance use, epilepsy), the mechanism of how rTMS induces neural plasticity remains poorly understood. Studying brain dynamics during stimulation could provide a crucial set of principles to optimize rTMS and other neuromodulation treatments (i.e., deep brain stimulation and vagus nerve stimulation).

The dynamics of neuronal plasticity are typically separated into the *induction* phase (changes during stimulation) and the *maintenance* phase (changes lasting minutes to hours after stimulation). The maintenance phase is characterized by a large change in synaptic response that persists for hours and involves modifications in gene expression, protein synthesis and synaptic function^1, 2, 3, 4, 5, 6^. The induction period has been less well-characterized, but in animal models changes observed during stimulation (induction phase) appear to relate to post-stimulation maintenance effects. During tetanic stimulation in rat hippocampal slices, synaptic responses dynamically changed as more pulses were applied^7^, and the magnitude of maintenance effects correlated with changes during stimulation. In non-human primates, >5Hz optogenetic stimulation to the sensorimotor cortices progressively increased functional connectivity during stimulation and predicted changes in post-stimulation connectivity ^8^. Furthermore, 10Hz magnetic stimulation (rTMS) to cat visual cortices elicited a pulse-wise increase in neural activity, resulting in significantly increased post-stimulation evoked and spontaneous activity ^9^. These findings in animal models suggest discrete neural changes occur during induction that relate to post-stimulation changes.

Few studies have examined the induction phase in humans. Two studies have recorded scalp electroencephalogram (EEG) while applying non-invasive rTMS and reported changes in the evoked potential during stimulation^10, 11^. However, due to the short latency of these evoked potentials and the possibility of residual stimulation-related artifacts, the interpretation of these findings is limited^12, 13^. In contrast, invasive recordings provide high spatiotemporal resolution with temporally defined artifact, allowing precise measurement of neural activity after each pulse. Recent studies using direct electrical stimulation coupled with invasive EEG demonstrated evidence of entrainment to the stimulation frequency^14^, decrease in low-frequency power during high frequency stimulation (100Hz)^15^, and increased theta activity directly after a stimulation train^16^. We recently demonstrated that repeated 10Hz electrical stimulation resulted in post-stimulation excitability changes in regions functionally connected to the stimulation site^17^. Furthermore, tracking the first pulse across stimulation trains was useful in predicting these post-stimulation changes, suggesting a potential link between the induction period and maintenance effects. Overall, human studies have begun to explore the complex dynamics of the induction phase of plasticity but a detailed characterization is still lacking.

In this investigation, we sought to better understand (1) the neural response to a stimulation train and (2) how brain activity during stimulation relates to post-stimulation connectivity changes. We measured cortical dynamics during repetitive stimulation across several cortical regions using invasive brain recordings in patients with medically-intractable epilepsy (Fig 1). We hypothesized that a stimulation train would increase neural responses in regions functionally connected to the stimulation site, and the strength of response during stimulation could be used to predict brain post-stimulation connectivity changes. During stimulation, we observed a consistent increase in high-frequency (70-170 Hz) power. Immediately following a train of stimulation, we observed an evoked response characterized primarily by low-frequency (1-8 Hz) power, which was highly correlated with the neural response during stimulation. Pre-stimulation connectivity as indexed by cortico-cortical evoked potentials and band coherence predicted the stimulation response. Further, in regions highly connected to the stimulation site, progressive change of the stimulation response was observed during the course of stimulation. Using features from the stimulation period, we were able to predict post-stimulation connectivity changes with high discriminability. This work demonstrates the feasibility of measuring neural activity during repetitive stimulation and serves to inform stimulation-based therapies for neuropsychiatric disorders.

**Figure 1.**
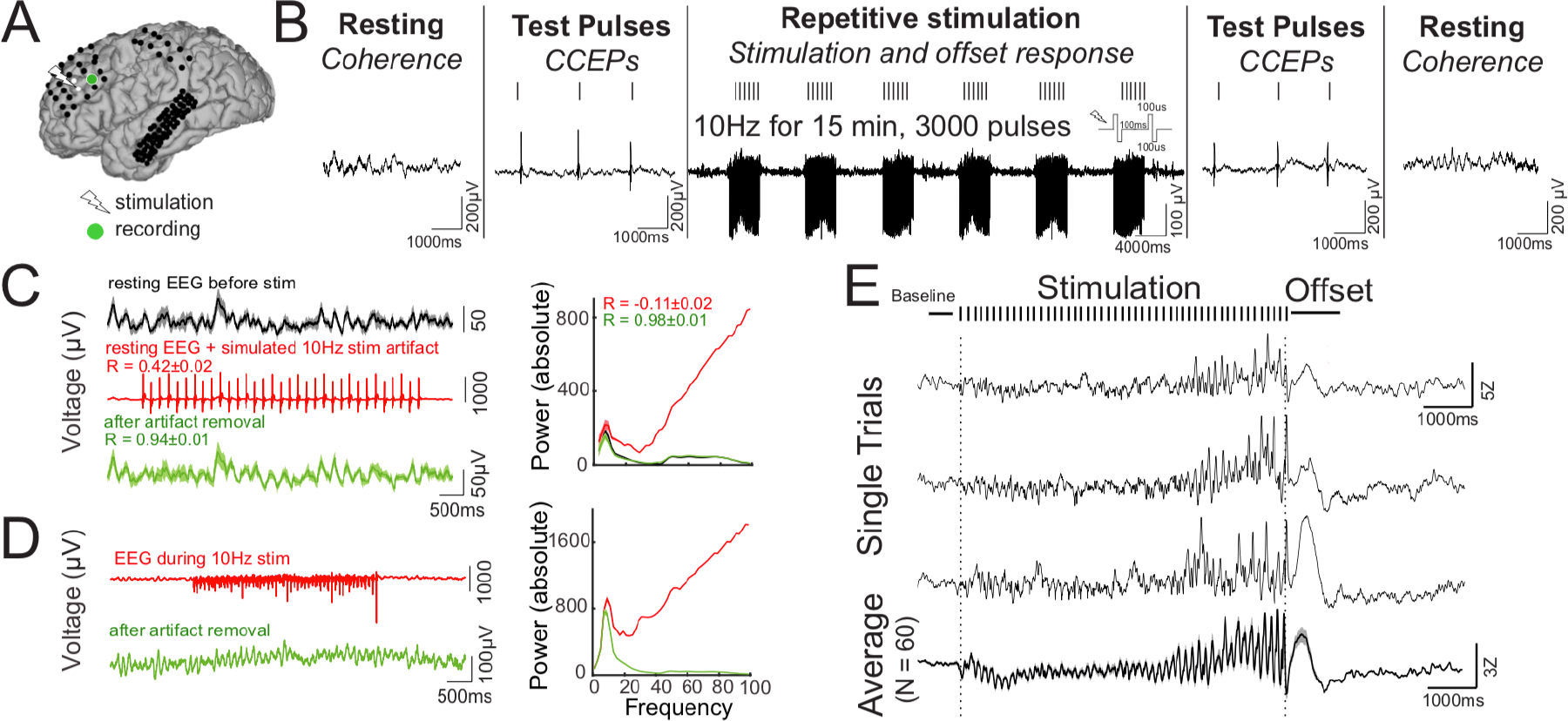
Experimental design and artifact removal. A) An example of intracranial stimulation and recording (Subject 4). Co-registered pre-operative MRI and post-operative CT allowed the visualization of subdural electrodes. B) Schematic of the stimulation paradigm. Periods of rest (‘Resting’) and single pulse cortico-cortical evoked potentials (CCEP, ‘Test pulses’) were applied before and after repetitive stimulation. Focal repetitive stimulation consisted of 15 minutes of 5s 10Hz trains with an inter-train interval of 6 to 10s. Pulses were bipolar with 100 μs/phase. Electrocorticography (ECoG) was recorded during each phase of the stimulation paradigm. C-D) Power-frequency analysis using real and simulated ECoG data to test the effectiveness of the artifact removal process. C) 10Hz stimulation artifact was added to the resting ECoG data, and the artifacts were subsequently removed using our artifact rejection pipeline (see Methods). The artifact-spiked data had poor correlation with the original data (R = 0.42±0.02) while the artifact-removed data had good fidelity with the original data (R= 0.94±0.01). Right panel: Power spectrum is showed for the original data, the artifact-spiked data, and the cleaned data. The power spectrum of the artifact-spike data is negatively correlated with that of the original data (R= −0.11±0.02), while the spectrum of the artifact-removed data resembled that of the original data (R= 0.98±0.01). D) Stimulation artifact removal applied to real ECoG during 10Hz stimulation. Right panel: Power spectrum before and after artifact removal. E) Single trial and average ECoG during and directly following the stimulation train recorded from the black electrode labeled in A. Note 1) the increase in evoked potential amplitude later in the stimulation train, and 2) the offset response: an evoked potential shortly after the 10 Hz stimulation train. Error bars show ±1 SEM.

## Methods

### Participants

Fourteen patients with medically-intractable epilepsy underwent surgical implantation of intracranial electrodes for seizure localization. Patient characteristics are described in Table 1. Patients were enrolled at two hospitals: North Shore University Hospital (Manhasset, New York, USA) and National Institute of Clinical Neurosciences (Budapest, Hungary). All patients provided informed consent as monitored by the local Institutional Review Board and in accordance with the ethical standards of the Declaration of Helsinki. The decision to implant, the electrode targets, and the duration of implantation were made entirely on clinical grounds without reference to this investigation. Patients were informed that participation in this study would not alter their clinical treatment, and that they could withdraw at any time without jeopardizing their clinical care.

**Table 1.** Participant characteristics, implant type and stimulation sites

| ID | Age | Gender | Handedness | Seizure Zone | Implant Type | Stimulation Location | MNI Coordinates |
| --- | --- | --- | --- | --- | --- | --- | --- |
| S1 | 21 | M | R | R mesial temporal | Grid/Strips | R precentral gyrus | 60, -12, 39 |
| S2 | 57 | F | L | R mesial temporal | sEEG | L precentral gyrus | -58, -6, 39 |
| S3 | 31 | F | R | R STG/mesial temporal | sEEG | R precentral gyrus | 57, -13, 37 |
| S4 | 43 | F | R | R posterior temporal | Grid/Strips | L middle frontal gyrus | -44, 35, 29 |
| S5 | 30 | F | R | L mesial temporal | Grid/Strips | L middle temporal gyrus | -35, 27, -29 |
| S6 | 32 | M | L | R middle frontal gyrus | Grid/Strips | R precentral gyrus | 65, -1, 19 |
| S7 | 20 | F | R | R frontal cortex | Grid/Strips | R precentral gyrus | 65, -5, 23 |
| S8 | 44 | F | R | R premotor cortex | sEEG | R middle frontal gyrus | 55, 28, 17 |
| S9 | 28 | F | R | R hippocampus | sEEG | R temporo-parieto-occipital junction | 51, -43, 21 |
| S10 | 28 | M | R | R middle frontal gyrus | sEEG | R inferior frontal gyrus | 19, 37, -21 |
| S11 | 36 | M | R | L temporo-polar-basal | sEEG | L mesial temporal | -23, 9, -32 |
| S12 | 50 | F | R | R OFC/amygdala | sEEG | R cingulate gyrus | 6, 37, 13 |
| S13 | 48 | F | R | R mesial temporal | sEEG | R middle frontal gyrus | 55, 35, 13 |
| S14 | 46 | M | R | R posterior temporal | sEEG | L inferior frontal gyrus | -51, 13, 4 |

### Electrode registration

Our electrode registration method has been described in detail previously^18, 19^. Briefly, in order to localize each electrode anatomically, subdural electrodes were identified on the post-implantation CT with BioImagesuite^4^, and were coregistered first with the post-implantation structural MRI and subsequently with the pre-implantation MRI to account for possible brain shift caused by electrode implantation and surgery^5^. Following automated coregistration, electrodes were snapped to the closest point on the reconstructed pial surface^6^ of the pre-implantation MRI in MATLAB^7^. Intraoperative photographs were previously used to corroborate this registration method based on the identification of major anatomical features. Automated cortical parcellations were used to localize electrodes to anatomical regions^8^.

### Electrophysiological recordings

Invasive electrocorticographic (ECoG) recording from implanted intracranial subdural grids, strips and/or depth electrodes were sampled at 512 or 2048Hz depending on clinical parameters at the participating hospital (U.S.A. and Hungary, respectively). Data preprocessing and analysis was performed using the FieldTrip toolbox^20^. Line noise (60Hz and 50Hz for recordings in U.S.A and Hungary, respectively) was removed using a notch filter. Direct electrical stimulation induced stereotyped stimulation artifacts that were ~10ms in duration. We applied a 4^th^ order bandpass filter (Butterworth, two-pass) in the 100-150 Hz frequency range, which sharply localizes the stimulation artifacts as these artifacts comprise primarily high frequency power (>40 Hz, Fig 1 C-D). Stimulation artifacts were subsequently detected by applying a value threshold. This value threshold was chosen per subject to detect all stimulation artifacts within the stimulation train. We replaced the stimulation artifact with stationary ECoG timeseries that represented similar amplitude and spectral profile as the background signal. This procedure was detailed previously^21^ and is preferred over simple spline interpolation given short intervals between pulses and the potential to introduce large spectral changes. To do this, we extracted ECoG signal with equal length as the stimulation artifact immediately preceding and following the artifact. We reversed the ECoG signal and applied a tapering matrix (1:1/n:0 for the preceding data, 0:1/n:1 for the following data, where n is the number of samples contained in the artifact). The two ECoG signals were added together and subsequently used to replace the artifact period. The effectiveness of this artifact removal process is shown in Figure 1. Following artifact rejection, we applied a bipolar montage to depth electrodes and a common average reference montage to grid electrodes^22^.

### Repetitive stimulation paradigm

In order to examine cortical dynamics during and after stimulation, we applied focal 10Hz stimulation in a clinically-relevant manner, as previously described^17^. Each subject received 15 minutes of 10Hz direct electrical stimulation in a bipolar fashion (biphasic pulses at 100 μs/phase). Each stimulation train was 5s (50 pulses / train) followed by 10s rest (15s duty cycle), resulting in 60 total trains and 3000 total pulses applied^10^ (Fig 1B). The stimulation current used was set at 100% of the motor threshold in patients with motor cortex coverage. Otherwise, 1 to 10mA was chosen depending on patient tolerance. The stimulation parameters were chosen to closely mimic commonly used rTMS treatment paradigms^11^. Stimulation sites were chosen in the frontal, temporal and parietal cortices as specified in Table 1 and Supplementary Figure 1.

### Temporal dynamics of the stimulation response

To examine cortical responses during the repetitive stimulation protocol both within the stimulation train (*intra-train*) and after the train (*post-train*), we epoched the ECoG signal surrounding the 5s stimulation train (3s before the first pulse and 8s after the last pulse). The epoch was subsequently standardized using Z-scores against the pre-train baseline period (−600ms to −100ms before the start of the first pulse). The *stimulation response* was defined as the mean response during the period of stimulation (first pulse to last pulse). The *offset response* was defined as the mean response from 10ms (to dissociate the offset response from the stimulation response) to 1000ms after the last pulse. To quantify the dynamics of the stimulation response over time, we used repeated-measures ANOVA. First, as evoked oscillations were prominent during the stimulation, we smoothed the broadband signal by applying a 4^th^ order Butterworth 1Hz low-pass filter. Second, we divided the stimulation period into 500ms bins. We reasoned if cortical excitability was changed during stimulation, then the means of individual bins should be significantly different. These bins represent the within-subject variable. Third, we created a variable representing the stimulation train number. Finally, we fitted a repeated measures model, where the broadband signal as stratified by the time bins is the response and the stimulation train number is the predictor variable (Time Bins ~ Stimulation Train). The F-statistic and associated p-value were obtained for each coefficient (Time Bins and Time Bins*Stimulation Train). The F-statistic for the Time Bins coefficient indicates if there was a significant effect of time during the stimulation on the broadband signal whereas the F-statistic for the Time Bins-Stimulation Train interaction indicates if there was progressive modulation of the broadband signal across the stimulation trains. A channel was considered stimulation responsive if either coefficient was significant using an p-value of 0.05 after FDR correction for multiple channels comparison.

### Spectral decomposition of intra-stimulation dynamics

We evaluated the time-frequency dynamics during stimulation using Hanning tapers (100ms interval, −1s pre-train to +2s post-train). We identified characteristic changes in spectral power of the stimulation train in three frequency ranges: 1-8Hz (low-frequency power), 8-40Hz (mid-frequency power), 70-170Hz (high-gamma power). These frequency bins were chosen after visual inspection of the time-frequency response due to observed differences in these bins (Figure 2A). To quantify slow changes in spectral power^23, 24^, we first applied a bandpass filter (Butterworth, two-pass) with filter order 4 for lower frequency bands (1-8Hz and 8-40Hz) or 8 for higher frequency bands (70-170Hz)^21^. Next, we took the absolute value of the filtered signal’s Hilbert transform to obtain the *analytic* signal (often referred to as band-limited power or BLP)^25^. Finally, we applied a 4^th^ order Butterworth 1Hz low-pass filter to obtain the slow component of the BLP ^24, 26^ in order to compare the different BLP signals. Each data point was Z-transformed relative to the pre-train baseline period (−600ms to −100ms before the start of the first pulse) for normalization across patients.

**Figure 2.**
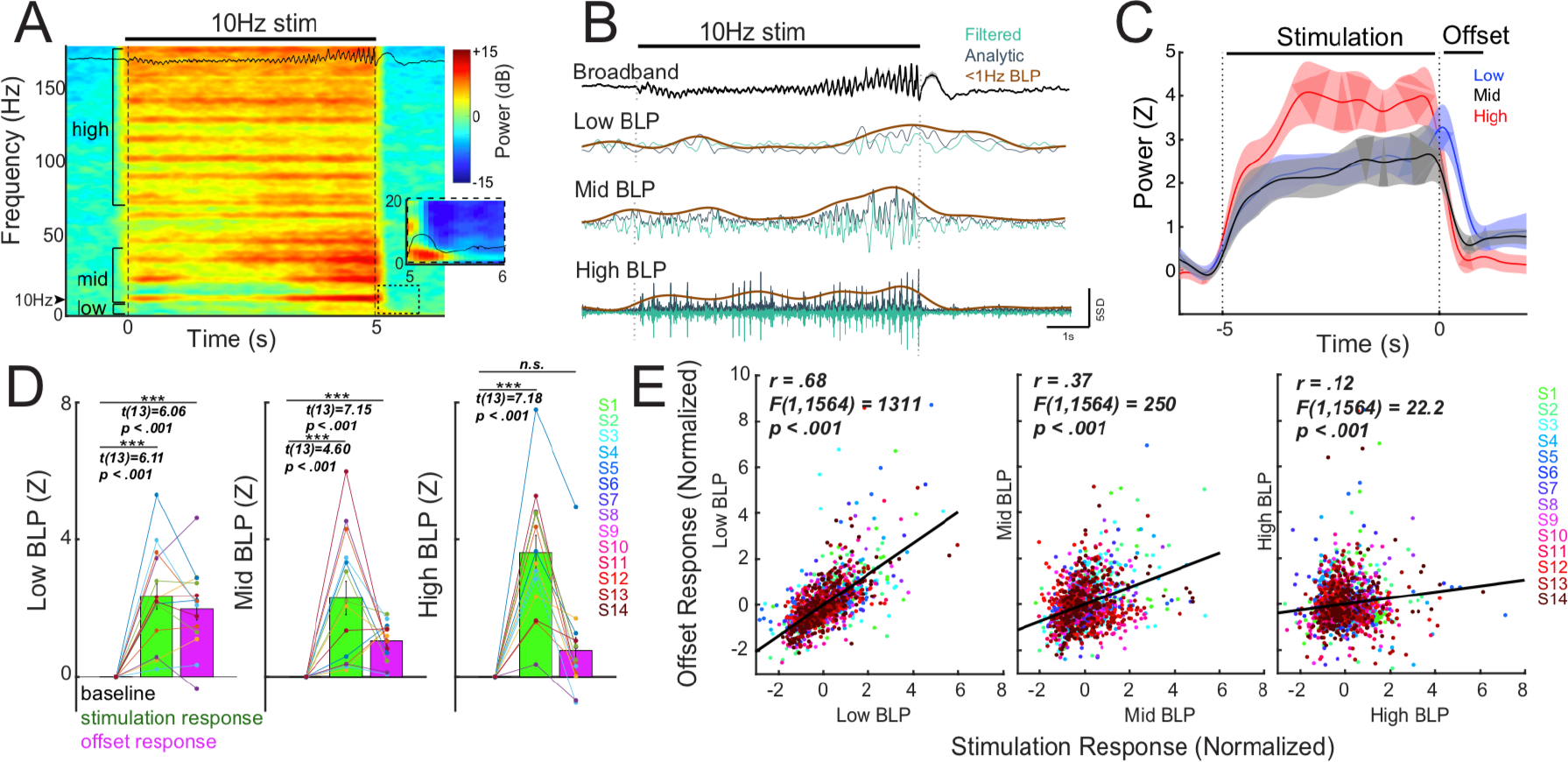
Repetitive stimulation elicits a multi-phasic neural response. A) Time frequency spectrogram during 10Hz stimulation. ECoG data shown in A-B are recorded from the electrode in Figure 1A. The stimulation period is characterized by changes in 8-40 Hz power, corresponding to the changes in evoked potentials observed in the raw broadband signal, as well as an increase in high gamma (>70 Hz) activity. Immediately after the stimulation train, an evoked response lasting ~1000ms occurs, and is primarily driven by low frequency power (<8Hz, see insert). B) Raw broadband signal (black) was transformed to band-limited power (BLP) to capture temporal dynamics of power changes during stimulation. C) Group dynamics of stimulation-induced response. Trials and channel data were averaged per subject, and the subject-averaged trace for each BLP is shown. Mid BLP (8-40 Hz) and 70-170 Hz power (high gamma power, HGP) remain elevated during stimulation and decrease quickly afterwards, while low BLP (1-8 Hz) increases during stimulation and peaks in the offset period. D) Group-level response during different phases of the stimulation train. Across subjects, the mean BLP during stimulation was significantly elevated during stimulation. In the offset period, low BLP and mid BLP was elevated, but not HGP. E) Channel-level relationship between the stimulation response and the offset response for each BLP with linear regression line (black line). Each color represents data from a single subject. Note the strongest correlation between the stimulation response and the offset response was observed with low BLP. Error bars show ±1 SEM; \*\*\**P* < 0.001.

### Pre/post-stimulation CCEP mapping (effective connectivity)

To examine causal changes in brain excitability at baseline and after stimulation, we performed cortico-cortical evoked potential (CCEP) mapping^27^. CCEPs have been used to predict the onset of ictal events^28^, examine the functional brain infrastructure^29, 30, 31, 32^, and causally examine the fronto-parietal^33^, hippocampal^34, 35^, visual^36^, and language^37^ networks. Prior to and immediately after repetitive stimulation, we applied bipolar electrical stimulation (biphasic pulses at 100 μs/phase) with a 1s inter-stimulation interval (ISI). This ISI was chosen to allow voltage deflections to return to baseline after ~500ms and to allow for sufficient trials to be collected within the expected time constraints in order to establish a stable pre-stimulation CCEP baseline^17^. A uniform random jitter (+/−200ms) was included in the ISI to avoid potential entrainment effects that could change baseline cortical excitability^17^. Stimulation current was chosen to match the current used during repetitive stimulation. 191 ± 20 (mean ± SE) single pulses were applied to assess the baseline CCEP while 662 ± 80 pulses were applied to assess post-stimulation CCEP. The number of CCEPs were chosen to maximize signal-to-noise within the amount of experimental time allotted. The number of pre and post CCEPs measured for each subject are outlined in Table 2. CCEPs from each channel were first epoched from −1000ms to 1000ms. The epoch was subsequently standardized using Z-scores against the pre-CCEP baseline period (−150ms to −50ms). The amplitude of the CCEPs was determined by averaging the standardized signal 10-100ms after the stimulation pulse. To evaluate whether CCEPs evoked at baseline were statistically different from zero, we used cluster-based nonparametric testing as previously described^38^. Briefly, we calculated one-sample t-statistics at every time point from 10ms to 100ms to form clusters of significant time points based on temporal adjacency at an alpha level of 0.05^38^. The cluster-level statistics were obtained by taking the sum of the t-statistic within each cluster. To generate the null distribution, we calculated the cluster t-statistic for randomly shuffled ECoG signals based on 1000 simulations and fitted a Gaussian curve. The cluster t-statistic was compared to this null distribution and the CCEP was considered significant using a p-value of 0.05 after FDR correction for multiple channels comparison. To determine if post-stimulation CCEPs were significantly different than baseline CCEPs (i.e. to assess presence of post-stimulation connectivity changes), we first matched the number of post-stimulation CCEP trials with baseline CCEP trials. For example, if 200 baseline CCEP trials were present, then the first 200 post-stimulation CCEP trials were used for statistical testing. From 10-100ms after the single pulse, two-sample t-statistic was obtained at every time point and significant clusters were formed based on temporal adjacency at an alpha level of 0.05. The sum of the t-statistic was obtained for each clusters. The null distribution for the cluster t-statistic was produced by randomly shuffling trials between baseline CCEPs and post-stimulation CCEPs for 1000 iterations and computing the cluster t-statistics. A Gaussian curve was fit over the null distribution. The post-stimulation CCEPs were considered significantly different from the baseline CCEPs using a cluster p-value of 0.05 corrected for multiple channels comparison. The percent of channels found to have significantly different post-stimulation CCEPs for each subject are outlined in Table 2.

**Table 2.** Stimulation setting, recording parameters and stimulation outcomes for each participant

| ID | Stimulation Frequency | Stimulation Current (mA) | Channel Number* | CCEP Number (pre-stimulation) | CCEP Number (post-stimulation) | Channels that are stimulation responsive (%)<sup>‡</sup> | Channel with significant response modulation (%)<sup>‡</sup> | Channel with pre/post CCEP change (%)<sup>‡</sup> | Channel with pre/post theta coherence change (%)<sup>§</sup> |
| --- | --- | --- | --- | --- | --- | --- | --- | --- | --- |
| S1 | 10Hz | 6 | 159 | 149 | 790 | 1.9 | 0.6 | 0 | 0 |
| S2 | 10Hz | 1.35 | 121 | 149 | 1265 | 18.2 | 7.4 | 0.8 | 0 |
| S3 | 10Hz | 1 | 166 | 144 | 330 | 0.6 | 0 | 0 | 0 |
| S4 | 10Hz | 8 | 104 | 194 | 794 | 42.3 | 19.2 | 7.6 | 7.7 |
| S5 | 10Hz | 7 | 176 | 231 | 868 | 12.5 | 1.1 | 5.1 | 0 |
| S6 | 10Hz | 10 | 57 | 149 | 448 | 63.2 | 28.1 | 50.9 | 0 |
| S7 | 10Hz | 10 | 62 | 149 | 448 | 53.2 | 27.4 | 16.1 | 3.2 |
| S8 | 10Hz | 5 | 26 | 149 | 448 | 26.9 | 0 | 15.4 | 0 |
| S9 | 10Hz | 5 | 64 | 149 | 743 | 21.9 | 1.6 | 0 | 0 |
| S10 | 10Hz | 10 | 79 | 149 | 449 | 31.7 | 6.3 | 3.8 | 0 |
| S11 | 10Hz | 8 | 54 | 149 | 297 | 77.8 | 20.4 | 18.5 | 0 |
| S12 | 10Hz | 6 | 99 | 199 | 400 | 19.2 | 8.1 | 3.0 | 0 |
| S13 | 10Hz | 4 | 197 | 358 | 997 | 18.8 | 7.6 | 0.5 | 1.5 |
| S14 | 10Hz | 4 | 202 | 358 | 1000 | 1.5 | 0.5 | 1.0 | 7.9 |

\*Depth bipolar channels used for analysis (stimulation channels and noise channels excluded)
<sup>‡</sup>The variations in these percentages across patients are dependent on the stimulation current. The correlation coefficient are as follows: Rcurrent-responsive: 0.70 (p=0.006); Rcurrent-modulation: 0.67 (p=0.009); Rcurrent-CCEP: 0.58 (p=0.030)
<sup>§</sup>The theta frequency band used is 4-8Hz with 29 out of 1566 total channels showing change. This analysis was performed for other frequency bands including 8-12Hz, 12-25Hz, 25-50Hz, and 70-100Hz. The numbers of channels showing change in these frequency bands were 30, 1, 3, 8 out of 1566 channels respectively.

### Pre/post-stimulation coherence analysis (functional connectivity)

To estimate functional connectivity through oscillatory synchrony of two brain regions, we computed coherence between all possible electrode pairs using FieldTrip (ft_connectivityanalysis)^39^. Coherence provides a measure of the phase difference between the paired signals that is unaffected by their amplitudes and has been previously used to estimate functional connectivity^8, 40^. To calculate coherence, we divided the pre and post-stimulation resting periods ranging from 5 to 10 minutes into 1s epochs. We used a multitaper method with 2 Hz spectral smoothing to compute the spectral estimate of each epoch^8, 41^. Coherence was calculated as the normalized cross-spectral density between the two signals. Theta coherence was obtained by averaging across frequency range of 4 to 8 Hz and was used in the primary analysis. Theta coherence has been previously used in both non-human primates and human studies to measure functional connectivity^8, 40^. Alpha (8-12Hz), beta (12-25Hz), gamma (25-50Hz) and high gamma (70-100Hz) frequency bands were also used for comparison analyses. To determine if pre-stimulation coherence for a particular pair of channels was significant, we generated a null distribution. To do this, we divided the resting timeseries into 20 bins, randomly shuffled the bins, and subsequently computed theta coherence. This bin number was chosen to be large enough to maintain the temporal structure of the ECoG time series but small enough to allow multiple iterations of data shuffling. This procedure was repeated for 1000 iterations as described above. A Gaussian curve was fit over the null distribution to yield the probability of observing a particular coherence value. Coherence for a particular pair of channels was considered significant using a p-value of 0.01 after FDR-correction for multiple channels comparison. To test for coherence differences pre and post-stimulation, we employed nonparametric testing as described previously^39^. Briefly, we calculated the Z-transformed coherence statistic and generated the null-distribution of the difference in coherence by randomly shuffling amongst pre and post-stimulation trials. This procedure was repeated 1000 times. A Gaussian curve was fit over the null distribution to yield the probability of observing the difference in coherence. Coherence was considered significantly different between two conditions using a p-value of 0.01 after FDR-correction for multiple channels comparison. The percent of channels found to have significantly different post-stimulation theta coherence (to the stimulation site) for each subject are outlined in Table 2.

### Prediction of post-stimulation connectivity change

To determine if features during the stimulation period predicted connectivity changes, we used logistical regression. For this analysis, we pooled all channels into a single dataset and categorized the channels by two outcomes: if there was significant pre/post CCEP change or significant pre/post theta coherence change. For a particular channel, the features derived from the stimulation period included (1) the stimulation response (the mean voltage during the stimulation period), (2) the presence of a significant stimulation response, and (3) the presence of modulation in the stimulation response after repeated stimulation trains. To evaluate the proportion of stimulation data required for good model performance, we created six subsets of the data. Using 1% (15 seconds), 10% (1.5 minutes), 20% (3 minutes), 60% (9 minutes), 85% (12.75 minutes), and 100% (15 minutes) of the stimulation trains, we derived the three features from the stimulation period. For each subset, we performed logistic regression with 10-fold cross validation. Receiver operating characteristic (ROC) curves were constructed and we quantified area under the curve (AUC) to evaluate model performance. To allow for comparison amongst models using different subset of data, we used bootstrap sampling (1000 permutations) to estimate the mean and 95% confidence interval of the model AUC.

## Results

We performed direct electrical stimulation using implanted electrodes while simultaneously recording electrical activity from the cortical surface (ECoG). Individual patient characteristics and stimulation sites are listed in Table 1. Electrode locations for each patient is visualized in Supplementary Figure 1. As shown in Figure 1A-B, our stimulation paradigm included resting periods to evaluate functional connectivity, test pulses to evaluate effective connectivity, and repetitive stimulation. This paradigm provided us the unique opportunity to assess neural activity before, during, and after stimulation.

### Repetitive stimulation elicits a characteristic neural response

In order to study plasticity induction, we sought to characterize neural activity occurring during repetitive stimulation. To do this, electrical stimulation artifacts must be carefully removed. We employed an artifact rejection procedure using principles previously validated for CCEPs^21^. To assess the validity of this procedure on stimulation artifacts during a stimulation train, we used a previously described simulation approach by adding stimulation artifacts to resting ECoG data^14^ (Fig 1C-D). Poor correlation was observed between the artifact-spiked data and the original data (R = 0.42±0.02) while good fidelity was observed between the artifact-removed data and the original data (Fig 1C, R= 0.94 ± 0.01). The artifact-containing data showed prominent high-frequency power (>40 Hz) while the resting data was mostly characterized by low-frequency power (<40 Hz). After artifact rejection, the power spectrum of the cleaned ECoG data resembled that of the original resting data (Fig 1C). Quantitatively, the power spectrum of the artifact-spike data is negatively correlated with that of the original data (R= −0.11 ± 0.02) while the spectrum of the artifact-removed data is highly correlated with that of the original data (R= 0.98 ± 0.01). At the exemplar channel outlined in Fig 1A, we demonstrated a consistent neural response to stimulation following artifact rejection, on both a single trial and average level (Fig 1E). This stimulation response consisted of an increasing evoked potential amplitude and a slow shift in voltage during the stimulation train. Immediately post-stimulation train, we observed an evoked response lasting roughly one second (offset response). We further characterized this signal by computing the time-frequency power spectrum (Fig 2A). High gamma power (HGP, 70-170 Hz) was elevated throughout the stimulation train, while 8-40 Hz power varied, which reflected the changing amplitude of the evoked potentials. The offset response showed primarily an increase in low-frequency power (1-8 Hz). To explore the temporal dynamics of these power changes during stimulation on coarse timescales, we computed slow changes in band-limited power (BLP, Fig 2B). HGP and mid-BLP (8-40 Hz) rapidly increased during stimulation and decreased quickly afterwards, whereas low BLP (1-8 Hz) gradually increased during stimulation and peaked during the offset period (Fig 2C). To quantify these changes, we averaged the response during stimulation (−5s to 0s) and during the offset period (0.01s to 1s). We found that stimulation elicited significant mean response in all BLP during stimulation (Fig 2D, paired t-test; low BLP: *t(13)* = *6.11*; mid BLP: *t(13)* = *4.60*; HGP: *t(13)* = *7.18*; all *P* < *0.001*). We then determined if a particular BLP was higher than the other during stimulation, and found HGP (70-170 Hz) to demonstrate a stronger response compared to low BLP (Supplementary Fig 2, left panel, paired t-test; *t(13)* = *3.50*, *P* = *0.003*) or mid BLP (*t(13)* = *3.29*, *P* = *0.006*). The offset period showed a significant increase in low BLP (Fig 2D, paired t-test; *t(13)* = *6.06*, *P* < *0.001*) and mid BLP (*t(13)* = *7.15*, *P* < *0.001*), but not HGP. Further, low BLP was significantly higher than both mid BLP (Supplementary Fig 2, right panel, paired t-test, *t(13)* = *3.30*, *P* = *0.006*) and HGP (*t(13)* = *3.52*, *P* = *0.004*) in the offset period.

We next asked if the offset response could be used as a proxy for the response during stimulation, as determining the response during stimulation in other modalities such as rTMS can be challenging due to the multitude of stimulation-related artifacts^12^. Thus, we evaluated the relationships between the stimulation response and the offset response. Linear regression was performed using data points from all patients. The strongest relationship was observed between the stimulation and the offset responses in low BLP (Fig 1E, *R* = *0.68*; F-test for the overall model, *F(1,1564)* = *1311*, *P* < *0.001*), followed by mid BLP (*R* = *0.37*, *F(1,1564)* = *250*, *P* < *0.001*). The weakest relationship was found in HGP (*R* = *0.12*; *F(1,1564)* = *22.2*, *P* < *0.001*). On a single trial level, the relationship between stimulation response and offset response was highest for low BLP (Supplementary Fig 3, left panel, *R* = *0.64*, *F(1,82518)* = *56079*, *P* < *0.001*), followed by mid BLP (Supplementary Fig 3, middle panel, *R* = *0.57*, *F(1,82518)* = *38868*, *P* < *0.001*) and HGP (Supplementary Fig 3, right panel, *R* = *0.51*, *F(1,82518)* = *28597*, *P* < *0.001*). Finally, we repeated the above analyses using raw voltage (broadband response) as this data would be easily accessible from a clinical perspective. Repetitive stimulation elicited a significant broadband response during stimulation (Supplementary Fig 4A, paired t-test; *t(13)* = *4.43*, *P* < *0.001*) and in the offset period (*t(13)* = *4.71*, *P* < *0.001*). A strong correlation was observed between the stimulation and offset response on both a channel (Supplementary Fig 4B, *R* = *0.45*, *F(1,1564)* = *370*, *P* < *0.001*) and single-trial level (Supplementary Fig 4C, *R* = *0.71*, *F(1,82518)* = *82676*, *P* < *0.001*).

**Figure 3.**
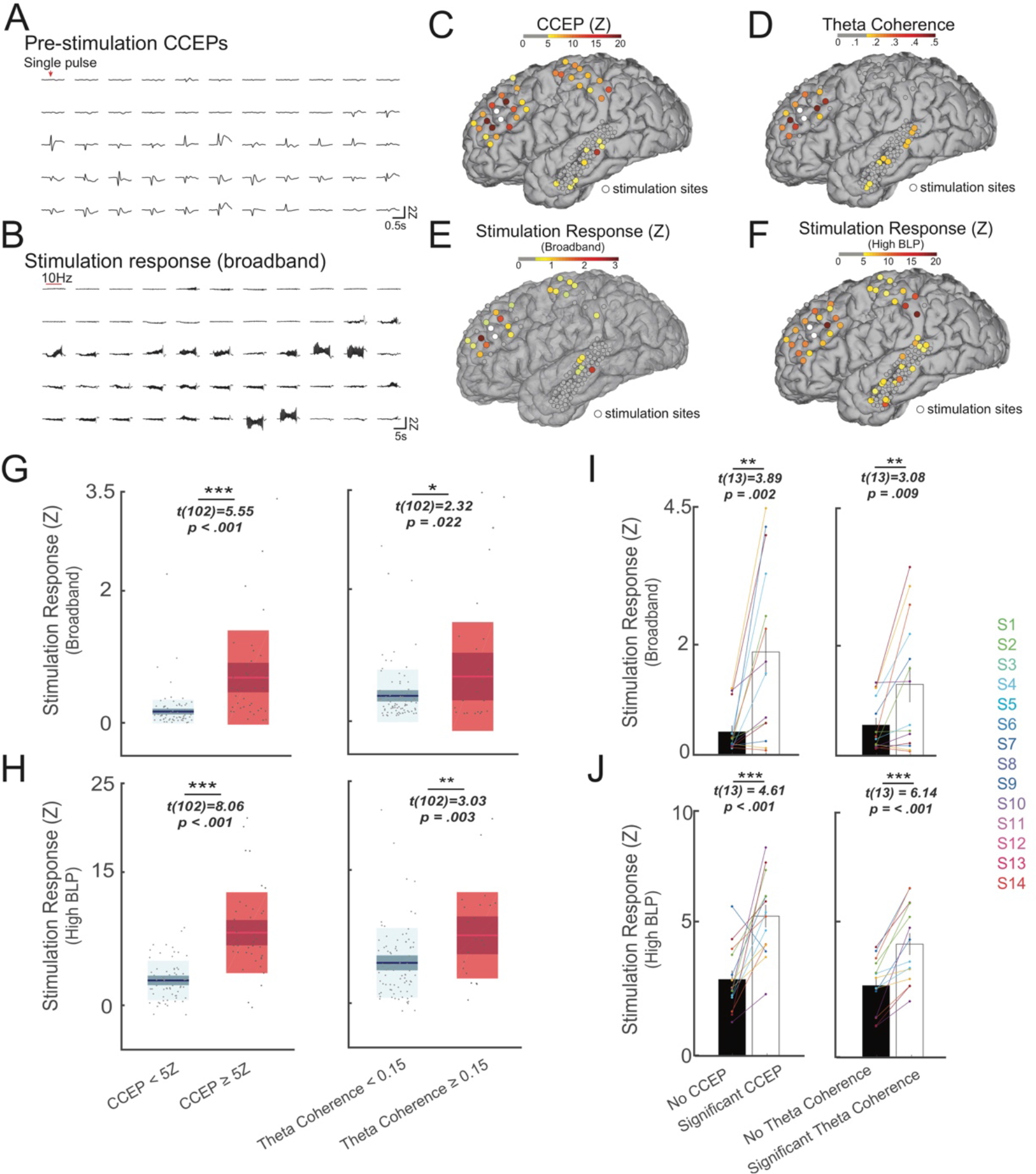
The stimulation response is predicted by functional and effective connectivity. A-B) Exemplar broadband signal across several channels for Subject 4. A) Single pulse cortico-cortical evoked potentials (CCEPs) recorded prior to stimulation and B) the corresponding neural response to the stimulation train. Qualitatively electrodes with strong CCEPs generally also elicited strong response during the stimulation train. C-F) Single subject (S4) spatial distribution of CCEP, theta coherence and the stimulation response (broadband and HGP). C) Strong CCEPs were elicited near the stimulation sites and at select parietal and temporal regions. D) Theta coherence to the stimulation site was highest in the prefrontal cortex. E) The mean voltage during stimulation (the broadband stimulation response) was high in certain regions across prefrontal, parietal and temporal cortices. F) Similar pattern of response was observed for HGP during stimulation. G) Increased mean broadband response was observed during stimulation in channels with higher pre-stimulation CCEP (left panel) and theta coherence to the stimulation site (right panel) H) Box plots demonstrating increased mean HGP during stimulation in channels with higher pre-stimulation CCEP (left panel) and theta coherence to the stimulation site (right panel). I-J) Channels with significant pre-stimulation CCEP or theta coherence were averaged per subject, and the mean stimulation response is shown for each subject. I) Higher broadband stimulation response is observed in channels with significant pre-stimulation CCEP (left panel) and theta coherence to the stimulation site (right panel) across subjects. J) Higher HGP stimulation response is observed in channels with significant pre-stimulation CCEP (left panel) and theta coherence to the stimulation site (right panel) across subjects. Error bars show ±1 SEM; \**P* < 0.05, \*\**P* < 0.01, \*\*\**P* < 0.001. Box plots show the mean value (innermost line), the 95% CI (dark band), and the SD (light band).

**Figure 4.**
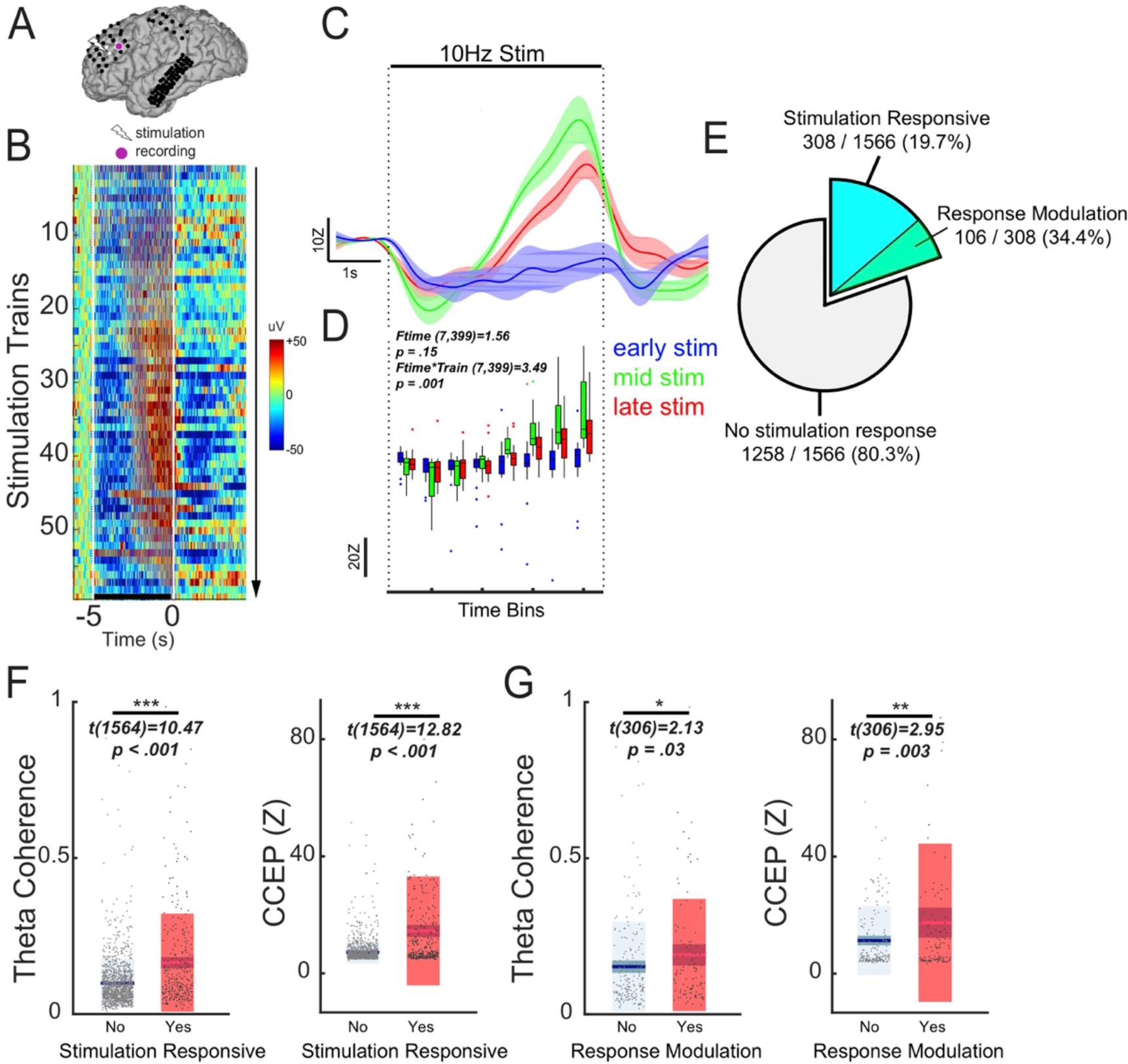
Progressive modulation of the stimulation response occurs in regions highly connected to the stimulation site. A-D) Example of neural changes across stimulation trains in one subject (S4). A) Location of stimulation and the recording electrode. B) Heatmap representation of the epoched broadband signal to increasing number of stimulation trains. Horizontal line represents time period of stimulation train. Color in the image represents the broadband signal during and after stimulation train. C) Time series of the smoothed broadband signal during stimulation as stratified by early (blue), middle (green), and late (red) trains in the stimulation protocol. D) Quantification of the stimulation dynamics in the exemplar channel. Repeated-measures ANOVA demonstrated a significant interaction between time during stimulation and the stimulation train number. E) Amongst aggregate of all channels across 14 patients, 308/1566 (19.7%) of channels were stimulation-responsive. Amongst the 308 stimulation-responsive channels, 106 (34.4%) showed a modulation in stimulation response (see Supplementary Fig 6 for location of these channels). F) Box plots stratifying pre-stimulation CCEP and theta coherence by stimulation responsive channels. Stimulation responsive channels demonstrated higher theta coherence (left panel) and CCEP amplitude (right panel). G) Box plots stratifying pre-stimulation CCEP and theta coherence by channels that did or did not undergo response modulation. Channels with stimulation response modulated by progressive trains demonstrated higher theta coherence (left panel, two-sample t-test, *P* = 0.03) and CCEP amplitude (right panel). Error bars show ±1 SEM; \**P* < 0.05, \*\**P* < 0.01, \*\*\**P* < 0.001. Box plots show the mean value (innermost line), the 95% CI (dark band), and the SD (light band).

In summary, repetitive stimulation elicits a measurable response both during and immediately after the stimulation train. The stimulation response is characterized by predominantly an increase in HGP whereas the offset response is driven primarily by low frequency power. The offset response is best correlated with the stimulation response using low frequency power; however, robust correlations were observed across all bands on a single trial level.

### Stimulation response is predicted by effective and functional connectivity

We next asked how the stimulation response relates to inter-area connectivity prior to stimulation. We quantified connectivity using single pulse CCEPs (a measure of network response to electrical stimulation) and resting theta coherence (a measure of related spontaneous neural activity). Qualitatively, channels with strong pre-stimulation CCEPs also exhibited strong broadband stimulation responses (Fig 3A-B) in one exemplar subject (S4). CCEPs were high locally at prefrontal electrodes near the stimulation site, as well as a subset of parietal and temporal cortices (Fig 3C). Pre-stimulation theta coherence between stimulation and recording electrodes was highest in prefrontal cortex (Fig 3D). During stimulation, the broadband response was high primarily in the prefrontal region (Fig 3E), whereas the HGP stimulation response was high in prefrontal and parietal cortices (Fig 3F). To relate pre-stimulation connectivity to the stimulation response, we plotted the stimulation response stratified by the strength of connectivity. In the same subject, we found that regions with stronger pre-stimulation CCEPs also demonstrated stronger broadband stimulation responses (Fig 3G, left panel, two-sample t-test; *t(102)* = *5.55*, *P* < *0.001*). Likewise, regions with higher pre-stimulation theta coherence also showed stronger broadband stimulation response (Fig 3G, right panel; *t(102)* = *2.32*, *P* = *0.022*). We repeated this analysis for the HGP stimulation response (Fig 3H), and found that HGP stimulation response was significantly higher in regions with higher pre-stimulation CCEPs (Fig 3H, left panel; *t(102)* = *8.06*, *P* < *0.001*) and theta coherence (Fig 3H, right panel; *t(102)* = *3.03*, *P* = *0.003*). To avoid using an arbitrary threshold and to generalize this finding across subjects, we next defined whether or not a particular channel showed significant baseline CCEPs or theta coherence to the stimulation site (see *Methods*). We found that across subjects the broadband stimulation response was consistently stronger in regions with significant pre-stimulation CCEPs (Fig 3I, left panel, paired t-test; *t(13)* = *3.89*, *P* = *0.002*) and theta coherence to the stimulation site (Fig 3I, right panel; *t(13)* = *3.08*, *P* = *0.009*). In a similar manner, the HGP during stimulation was higher in regions with significant pre-stimulation CCEPs (Fig 3J, left panel; *t(13)* = *4.61*, *P* < *0.001*) and theta coherence to the stimulation site (Fig 3J, right panel; *t(13)* = *6.14*, *P* < *0.001*). To test if these results were dependent on the coherence band used, we computed coherence using standard frequency bands (Supplementary Fig 5). Channels with significant band coherence in all frequency bands showed stronger broadband stimulation response (paired t-test, all P < 0.05). Higher HGP stimulation response was observed in channels with significant pre-stimulation band coherence across alpha, beta and gamma frequency (all P < 0.01), but not high gamma frequency (P = 0.08). In summary, pre-stimulation network connectivity defined by CCEPs and coherence across frequency bands predicts the neural response during stimulation.

**Figure 5.**
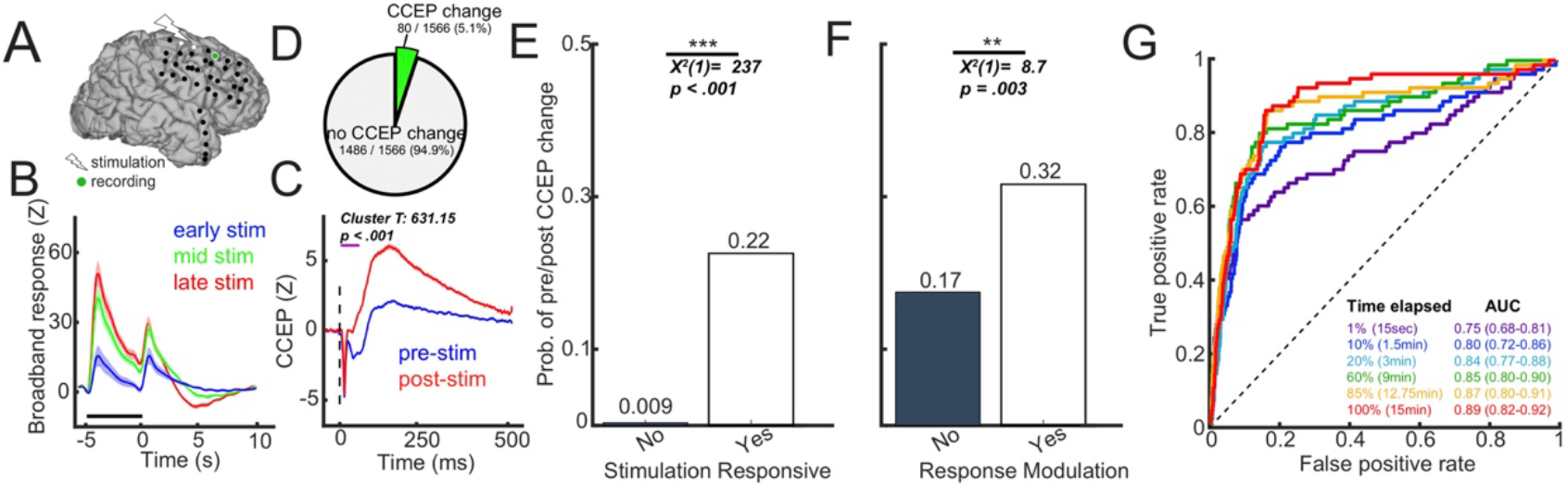
The stimulation period predicts connectivity changes following the entire stimulation protocol. A-C) Exemplar channel recording from Subject 5. A) Location of stimulation and the recording electrode. B) Time series of the smoothed broadband signal during stimulation as stratified by early (blue), middle (green), and late (red) trains in the stimulation protocol. C) The corresponding pre/post single pulse CCEP. Note the similar direction of change in CCEP and in the stimulation response. D) Amongst aggregate of all channels across 14 patients, 80/1566 (5.1%) of channels showed significant pre/post CCEP change. E) The probability of CCEP change in regions with and without a significant stimulation response. Stimulation responsive channels had higher probability of showing pre/post CCEP change. F) The probability of CCEP change in regions with and without response modulation by stimulation trains. Amongst only the stimulation-responsive channels, those that showed response modulation had a higher probability of showing pre/post CCEP change. G) Receiver operating characteristic (ROC) curves using features from the stimulation period to predict pre/post CCEP change. The features used were the presence of significant stimulation response without response modulation, the presence of response modulation by stimulation trains and the mean amplitude of the broadband signal during stimulation. Note aside from using 1% of the stimulation protocol, model performance was similar using different amount of the stimulation data. Error bars show ±1 SEM.

### Stimulation response is modulated after repeated stimulation in regions highly connected to the stimulation sites

As shown in Figure 1B, our stimulation paradigm included 60 repeated applications of 10Hz stimulation trains separated by resting periods. This allowed us to track changes in neural activity during each stimulation train. At the exemplar channel from Figure 1A, we observed that the broadband signal during stimulation starts changing around train 10 and peaks in amplitude around train 40 (Fig 4B, 4C). Compared to early stimulation, later stimulations elicited progressively more negative response at the beginning of the train, and more positive response towards the end of the train (Fig 4B, 4C). To quantify the stimulation dynamic, we used a repeated-measures model incorporating time bins and stimulation train as variables (see Methods and Figure 4D). A significant interaction between time during stimulation and the stimulation train number (Ftime*train(7,399) = 3.49, P = 0.001) indicated that there was modulation of the stimulation response over repeated trains. Across all channels, 19.7% of channels (308/1566) were stimulation responsive, which is defined as significance of either the time or time*train coefficients in the repeated-measures model (Fig 4E). Within these stimulation responsive channels, 34.4% (106/308) showed progressive modulation of the stimulation response. Spatially, these stimulation responsive channels were primarily local to the stimulation site (Supplementary Fig 6). On a single subject level, the stimulation amplitude correlated strongly both with the number of stimulation responsive channels (R = 0.70, P = 0.006, Table 2) and the proportion of channels exhibiting response modulation (R = 0.67, P = 0.009). Pooling together all channels, stimulation responsive channels exhibited higher pre-stimulation theta coherence (Fig 4F, left panel, two-sample t-test; *t(1564)* = *10.47*, *P* < *0.001*) and CCEP (Fig 4F, right panel; *t(1564)* = *12.82*, *P* < *0.001*) compared to non-responsive channels. Furthermore, limiting the analysis to only the stimulation responsive channels (N = 308), those channels which demonstrated modulation of the stimulation response over time had higher pre-stimulation theta coherence (Fig 4G, left panel, two-sample t-test; *t(306)* = *2.13*, *P* = *0.03*) and CCEP (Fig 4G, right panel; *t(306)* = *2.95*, *P* = *0.003*) compared to channels that were stimulation responsive but did not undergo modulation of activity across stimulation trains. In summary, stimulation-responsive regions were more strongly connected to the stimulation site than non-responsive regions. In a subset of regions that were strongly connected to the stimulation site, repeated stimulation trains progressively modulated neural activity over time.

### The stimulation period predicts changes in post-stimulation connectivity

Finally, we asked if features pertaining to the stimulation period can be used to predict post-stimulation connectivity changes. We hypothesized that stimulation responsive regions and regions which exhibit modulation of the stimulation response would be more likely to show post-stimulation connectivity changes. In an exemplar channel (Subject 5), single 10Hz stimulation trains elicited a strong stimulation response, and repeated trains elicited progressive modulation of the response (Fig 5A-B). We also observed a significant increase in the CCEP after the entire repetitive stimulation protocol (Fig 5C; non-parametric cluster T-test, cluster T = 631.15, *P* < *0.001*). Of note, the direction of amplitude change observed in the pre/post CCEP (here, stronger post-stimulation CCEP) mirrored that of the direction of modulation (stronger) of the broadband stimulation response (Fig 5B-C). In total, a small proportion (5.1%, 80/1566) of all channels showed a significant pre/post CCEP change (Fig 5D). Likewise, a small proportion (1.9%, 29/1566) of all channels demonstrated significant change in pre/post theta coherence (Supplementary Fig 7A). We found that stimulation responsive channels were more likely to undergo pre/post CCEP change (Fig 5E, 22.0% vs 0.90%, chi-squared test, *χ*^2^*(1)* = *237*, *P* < *0.001*). Similarly, of only the stimulation responsive channels, channels where stimulation response was progressively modulated were more likely to show pre/post CCEP change (Fig 5F, 32.0% vs 17.0%; *χ*^2^ = *8.7*, *P* = *0.003*). Additionally, we performed this analysis looking at post-stimulation changes in theta coherence. The presence of a significant stimulation response was also associated with post-stimulation change in theta coherence (Supplementary Fig 7B, 3.3% vs 1.5%; *χ*^2^ = *4.1*, *P* = *0.042*). The presence of response modulation trended towards higher probability of pre/post theta coherence change (Supplementary Fig 7C, 5.7% vs 2.0%; *χ*^2^ = *3.0*, *p* = *0.083*).

We next used logistic regression to assess if post-stimulation connectivity changes can be predicted from features during stimulation. The features used were (1) the presence of a significant stimulation response without response modulation (binary variable), (2) the presence of response modulation (binary), and (3) amplitude of the broadband stimulation response (continuous). We used subsets of this data to determine the minimal number of stimulation trains required for stable model performance. Using 1%, 10%, 20%, 60%, 85% and 100% of the stimulation data, post-stimulation changes in CCEP were predicted with AUC 0.75 (95% confidence interval: 0.67-0.81), 0.80 (0.72-0.86), 0.84 (0.77-0.88), 0.85 (0.80-0.90), 0.87 (0.80-0.91), 0.89 (0.82-0.92), respectively (Fig 5G). Aside from the 1% data subset, the AUC for all other subsets were not significantly different, as the confidence intervals were overlapping. We were not able to meaningfully predict post-stimulation changes in theta coherence (Supplementary Fig 7D). In summary, changes in CCEP and theta coherence occurred in a small proportion of total channels. Features from the stimulation period predicted CCEP changes after the stimulation protocol with high discriminability. Using subsets of the stimulation trains, we showed that model performance can reach stability with only a small proportion of the total data.

## Discussion

In this study we investigated the neural dynamics during and after a series of repeated stimulation trains. Across several stimulation sites, we observed a consistent increase in HGP activity during the stimulation train and a slower post-train evoked response that strongly correlated with activity changes during the train. We showed that in areas highly connected to the stimulation site, as indexed by two measures of connectivity, the stimulation response was stronger and exhibited progressive modulation with repeated trains. Finally, the stimulation period predicted post-stimulation connectivity changes with high discriminability. Importantly, we demonstrated that using a subset of the stimulation protocol was sufficient for good model performance.

Mounting evidence suggests that the induction period is characterized by stimulation-driven cycles of excitation and inhibition. In this study, we expand on evidence from animal studies during the induction period and offered insight into how a clinically-relevant stimulation pattern influences brain dynamics. First, we observed an increase in HGP during stimulation, especially in regions functionally connected to the stimulation site (Fig 2-3). As HGP has been shown to correlate with spiking activity^26, 42, 43^, this work suggests that 10Hz stimulation elicits increases in neuronal activity during stimulation. These findings are similar to recent work in rat hippocampal slices which reported a slow voltage drift during high frequency stimulation that corresponded with the change in excitatory post-synaptic potential (EPSP) amplitude^7^. Second, low-frequency power also increased during 10Hz stimulation. Although this is in contrast with a recent study that reported stimulation-driven decreases^15^, this may be attributed to the difference in frequency of stimulation (10Hz vs 100Hz). Finally, after each stimulation train, we observed an evoked potential lasting for roughly a second, which we termed the offset response. Consistent with a prior study, this offset response is primarily driven by low-frequency power not be dependent on stimulation frequency^40^. These slow waves likely represent GABA-ergic inhibitory periods^44^, which have been observed during spike and wave discharges^45, 46^ as well as single pulse evoked potentials in animals^47^ and humans^27, 45^. Together, these findings suggest that in regions functionally connected to the stimulation site, stimulation trains increased spiking activity, which were followed by an inhibitory rebound period.

Furthermore, in regions highly connected to the stimulation site, we observed a modulation of the stimulation response as multiple trains were applied (Fig 4). This may represent reorganization of functional networks during stimulation^8^. In non-human primates, repetitive optogenetic stimulation was able to strengthen functional connectivity between motor and somatosensory cortices within minutes. In the same study, stimulation-evoked activity predicted post-stimulation changes in connectivity. To our knowledge, we demonstrated this phenomenon in the human cortex for the first time. Specifically, we found that post-stimulation connectivity changes were more likely to occur in regions that showed progressive modulation of the stimulation response (Fig 5). To corroborate this, we demonstrated that the stimulation period predicted post-stimulation CCEP changes. Of note, although we were able to predict post-stimulation CCEP changes with high discriminability, our model performance was not as robust in the prediction of post-stimulation theta coherence changes. This could be due to the order of post-stimulation testing (resting data for coherence was collected after CCEP testing and therefore was further in time from the stimulation protocol) or to the method of connectivity measurement (evoked for CCEPs, rest for coherence analysis). Although it was beyond the scope of this investigation, comparing theta coherence and CCEP in approximating network connectivity would be of substantial interest.

The clear neurophysiological effects we observed during stimulation offer intriguing clinical utility. Efforts to optimize treatment by updating stimulation parameters in real-time have recently sparked an interest in closed-loop brain stimulation^48, 49, 50^. Two findings from this study highlight the potential for real-time implementation. First, we found that the strength of stimulation response was correlated strongly with the strength of the offset response, both on a channel and single trial level. This is an important finding given removal of stimulation artifacts in other modalities (i.e. EEG during rTMS^51^) is often difficult. Our findings suggest that the offset response may be used as a proxy for the stimulation response during clinical monitoring. Second, we found that only a few minutes of stimulation data were required to achieve stable model performance in predicting post-stimulation effects. To date, we lack a method to rapidly optimize stimulation patterns for an individual, as pre/post testing after the entire stimulation protocol will be cumbersome if multiple parameters are tested. Yet based on our results, if the stimulation response (or offset response) can be monitored and pre-processed in real-time or near real-time, then effects of multiple stimulation paradigms can be tested rapidly. If this work can be replicated by non-invasive modalities such as TMS, then one could envision a ‘stimulation localizer’ day prior to stimulation treatment. During this day, pre-stimulation characteristics would first be used to help localize the stimulation site (and network) of interest^17^. This would be followed by multiple stimulation trains of short duration (2 minutes) with various parameters (frequency, pattern, intensity) to select the stimulation paradigm that would maximize post-stimulation effects.

Several aspects of this study limit its generalizability. First, since seizures can alter both local and global brain excitability and connectivity^52, 53, 54^, our results may not be entirely representative of responses in a healthy brain. Although direct recording provides unsurpassed spatiotemporal resolution in humans, epilepsy patients differ in their underlying etiologies and electrode implantation patterns. Second, behavioral effects of stimulation were not measured in this study and this warrants further investigation with mood self-reporting^55^. Finally, given hospital time constraints (typically ~1 hour per patient), we were unable to modify the stimulation parameters that may be critical in inducing plasticity: stimulation site, frequency, and intensity.

Future work includes (1) a thorough examination of how parameters of a *single stimulation train* – including train frequency, number of pulses, and intensity – affect stimulation and offset responses; (2) an evaluation of how both induction and post-stimulation effects are modulated when repeated trains (3000 pulses each) are applied at different stimulation frequencies (10 and 100Hz for example), stimulation intensities (50% and 100% MT), and stimulation sites; (3) an evaluation of not only how these stimulation parameters affect mood, similar to Chang et al.^15^, but also for several stimulation frequencies and targets; (4) replication of these experiments using microelectrode recording in non-human primates to evaluate mechanistic changes at a scale unattainable even in ECoG; (5) adapting these experiments to patients with neuropsychiatric disorders using non-invasive neuromodulation (TMS) paired with scalp EEG, in order to determine if these signals can be feasibly measured and monitored for clinical translation. Knowledge gained from these planned experiments will greatly enhance our understanding of how stimulation modulates human brain activity and behavior, helping propel us to the next generation of personalized neuromodulation therapies.

## Conclusions

Here, we characterized the neural activity in the time period surrounding a train of electrical stimulation and its dependency on existing functional networks, thus providing valuable insight as to how the brain changes during stimulation. Furthermore, we demonstrated the utility of this information, by showing that neural activity during stimulation serves to predict stimulation-induced changes in network connectivity. Taken together, our work provides key insights into the development of closed-loop neuro-modulatory devices.

## Acknowledgements

The authors thank Maria Fini and Victor Du for help with data collection; Pierre Megevand and Erin Yeagle for help with technical considerations of the experimental design; Amit Etkin, and Wei Wu for comments on the manuscript. All authors discussed the data, analysis and methods and contributed to the manuscript. The authors are enormously indebted to the patients that participated in this study, as well as the nursing and physician staff at North Shore University Hospital (Manhassat, NY) and the National Institute of Clinical Neurosciences (Budapest, Hungary). The authors declare no competing financial interests. C.J.K was funded by the National Institute of Neurological Disorders and Stroke (F31NS080357-01 and T32-GM007288), Stanford Society of Physician Scholars Collaborative Research Fellowship, and Alpha Omega Alpha Postgraduate Research Award. D.F. was funded by the Hungarian National Research, Development and Innovation Office (2017-1.2.1-NKP-2017-00002).

## Authorship

Y.H., and C.J.K performed data analysis and data visualization; C.J.K and A.D.M. were involved in the design and conception of the research project; C.J.K., B.H., L.E., D.F., and J.L.H., acquired the data; C.J.K., and Y.H., wrote the manuscript; all authors were involved in critical reading and revision of the manuscript.

## Competing interests

The authors declare no competing financial interests.

