## Supplemental Information for "Intracortical dynamics underlying repetitive stimulation predicts changes in network connectivity"

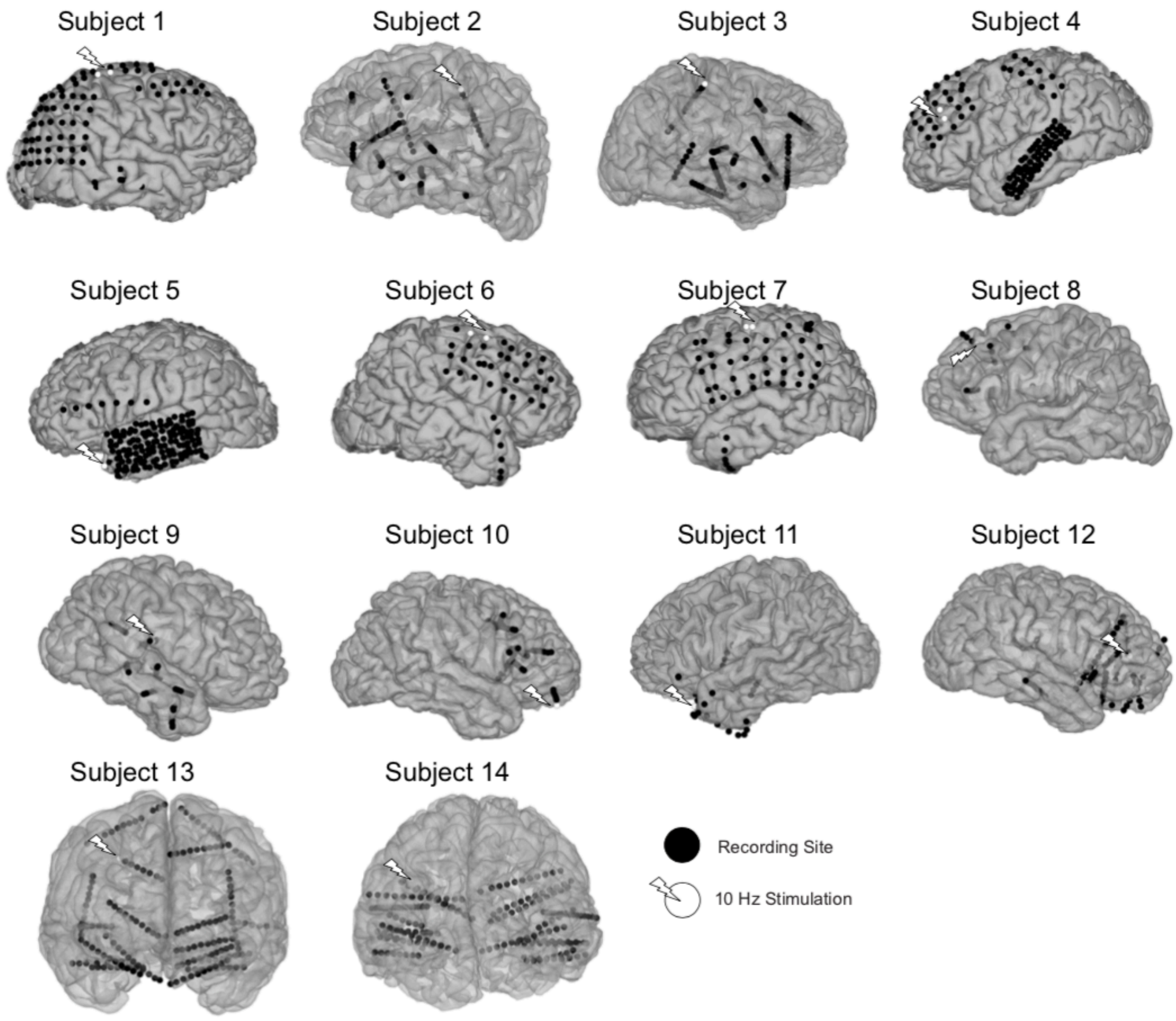


**Supplementary Figure 1 – Stimulation sites and electrode coverage.** Single-subject brain plots visualizing the location of the stimulation sites and the recording electrodes. White electrodes indicate the stimulation sites while black electrodes indicate recording channels. For each subject, 15 minutes of 10Hz focal direct electrical stimulation was applied in a bipolar fashion (see Methods).

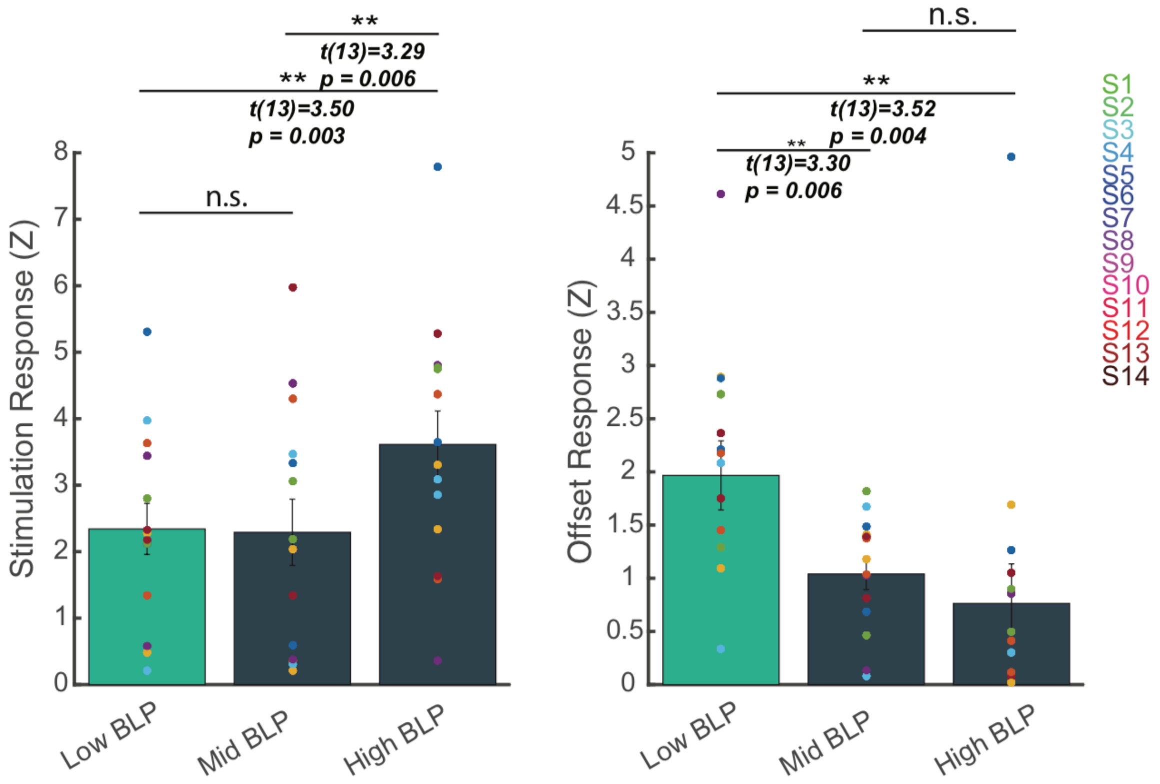


**Supplementary Figure 2 – Comparison of the stimulation response and the offset response as stratified by band-limited power.** During stimulation, the mean HGP was significantly higher than low BLP (paired t-test, $P = 0.003$) and mid BLP ($P = 0.006$). In the offset period, low BLP was significantly higher than the mid BLP ($P = 0.006$) and HGP ($P = 0.004$). Each dot represents the channel-averaged response for a single subject. Error bars show ±1 SEM; $^{**}P < 0.01$, $^{***}P < 0.001$.

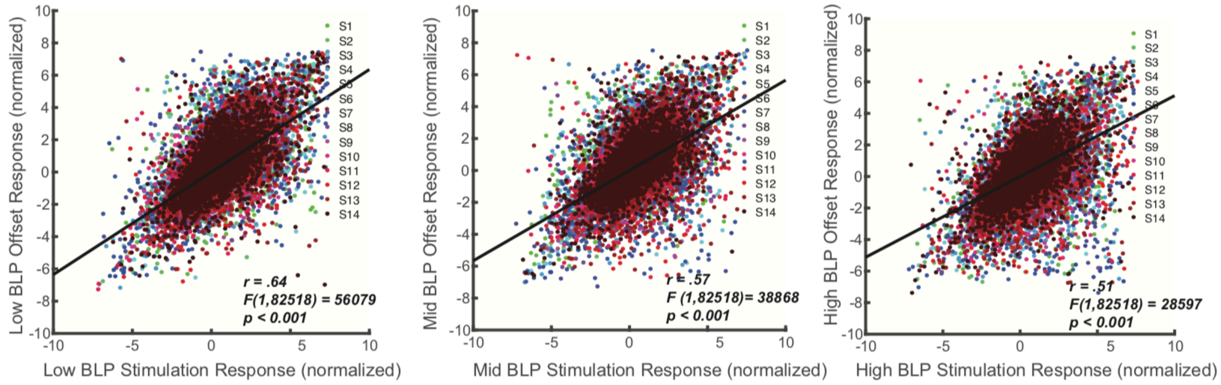


**Supplementary Figure 3 – The stimulation response is highly correlated with the offset response on a single trial level.** For each individual stimulation train, the response during the stimulation period was correlated with the response during the offset period. This was done for every channel per subject, generating (number of trains * number of channels) data points for each subject. The linear regression line (black) was calculated using the aggregate of all data points. In all 3 BLP, the stimulation response was highly correlated with the offset response on a single trial basis.

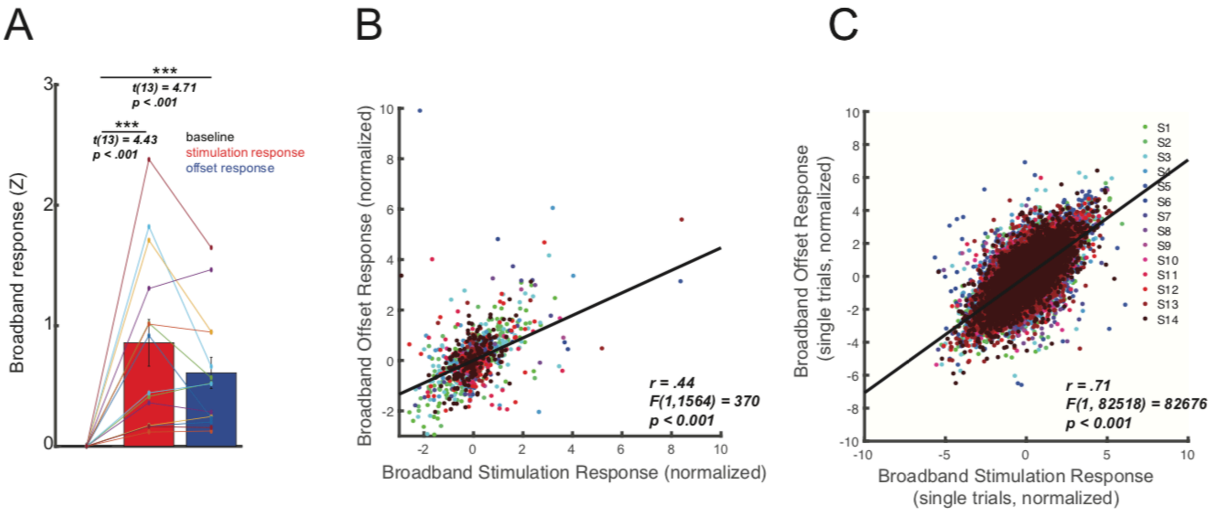


**Supplementary Figure 4 – Quantification and correlation of the stimulation and offset responses using broadband ECoG signal.** A) Repetitive stimulation elicited a significant broadband response during stimulation (paired t-test, $P < 0.001$) and a significant broadband response in the offset period ($P < 0.001$). Each dot represents the channel-averaged response per subject. B) The broadband stimulation response is correlated with the broadband offset response. Each dot represents a single channel response. Linear regression is performed over all data points (black line). C) On a single trial level, the broadband stimulation response is correlated with the broadband offset response. Each dot presents a single trial. Linear regression is performed over all data points (black line). Error bars show ±1 SEM; ***$P < 0.001$.

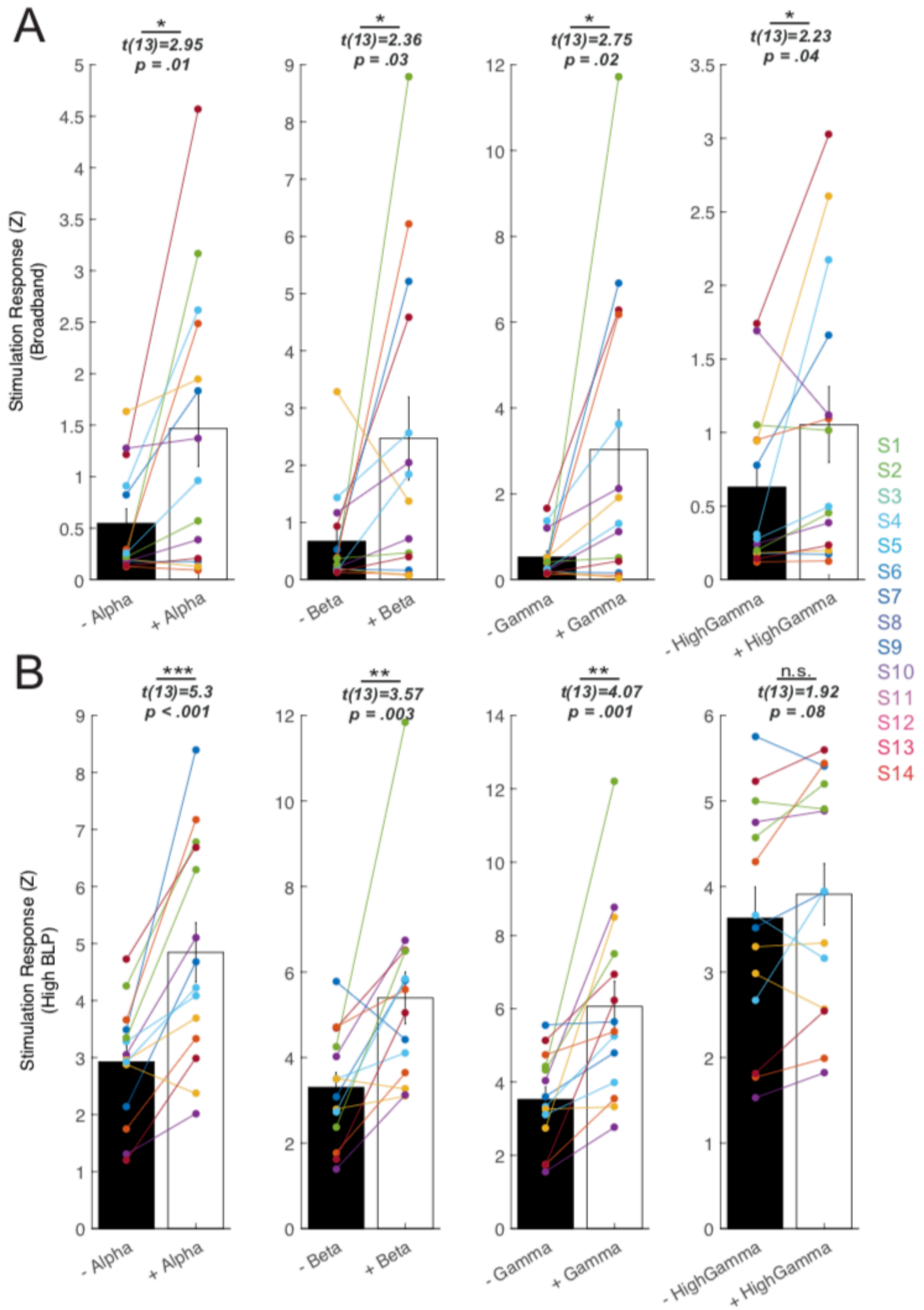


**Supplementary Figure 5. Stimulation responses as stratified by multiple coherence bands.** A-B) Channels with significant pre-stimulation band coherence were averaged per subject, and the mean stimulation response is shown for each subject. A) Higher broadband stimulation response is observed in channels with significant pre-stimulation band coherence across alpha (8-12Hz), beta (12-25Hz), gamma (25-50Hz) and high gamma (70-100Hz) frequency (paired t-test, all $P < 0.05$). B) Higher HGP stimulation response is observed in channels with significant pre-stimulation band coherence across alpha (8-12Hz), beta (12-25Hz) and gamma (25-50Hz) frequency (all $P < 0.01$), but not high gamma (70-100Hz) frequency ($P = 0.08$). Error bars show ±1 SEM; *$P < 0.05$, **$P < 0.01$, ***$P < 0.001$.

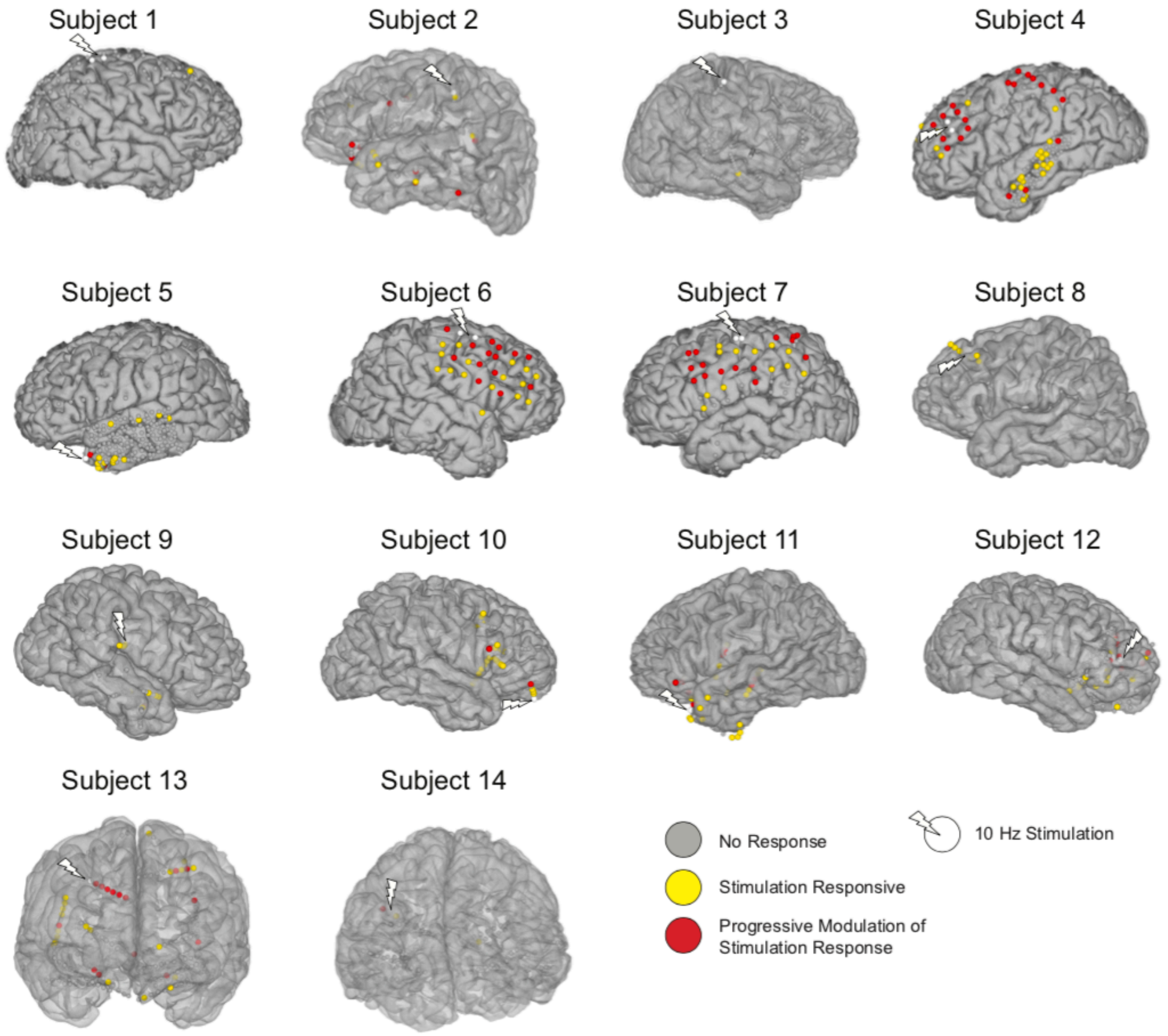


**Supplementary Figure 6 – Location of electrodes exhibiting significant stimulation responses and modulation of the stimulation response.** In each subject, significant stimulation responses were elicited in a subset of the regions recorded (yellow and red electrodes). Of all the stimulation-responsive electrodes, some underwent significant change in the stimulation response over repeated application of the stimulation train (red electrodes). Regions demonstrating response modulation are primarily local to the stimulation sites, although a small proportion of these electrodes are found on the opposite hemisphere or distant cortex.

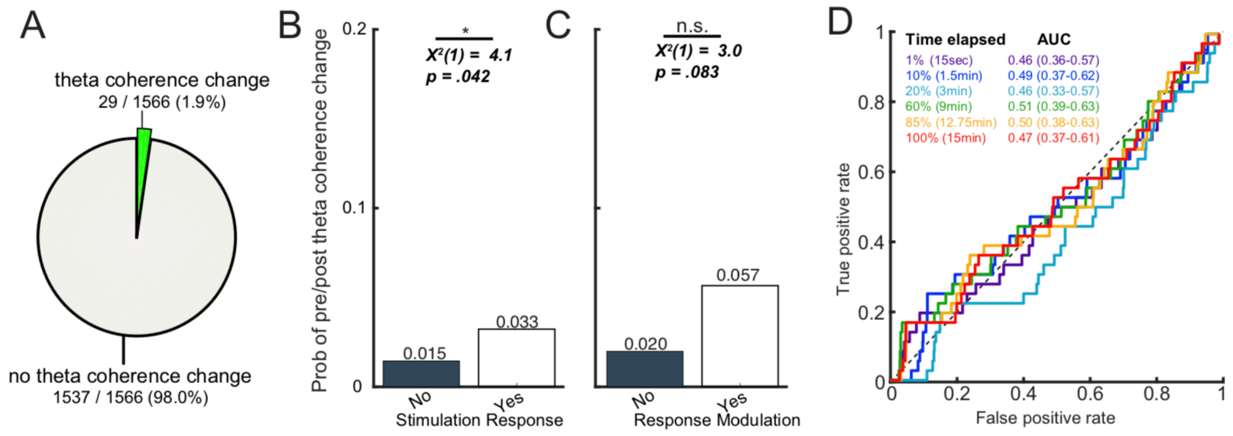


**Supplementary Figure 7 – Prediction of post-stimulation theta coherence change using features from the stimulation period.** A) Amongst aggregate of all channels across 14 patients, 29/1566 (1.9%) of channels showed significant pre/post theta coherence change. B) The probability of theta coherence change in regions with and without significant stimulation response. Regions exhibiting a significant stimulation response had higher probability of showing pre/post theta coherence change (chi-squared test, *P = 0.042*). C) The probability of theta coherence change in regions with and without modulation of the stimulation response. Amongst all stimulation-responsive channels, those that show response modulation trended towards a higher probability of showing pre/post theta coherence change (*P = 0.083*). C) Receiver operating characteristic (ROC) curves using features from the stimulation period to predict pre/post theta coherence change. The features used were the presence of significant stimulation response without response modulation, the presence of response modulation by stimulation trains and the mean amplitude of the broadband signal during stimulation.
